# Psilocybin prevents habituation to familiar stimuli and preserves sensitivity to sound following repeated stimulation in mouse primary auditory cortex

**DOI:** 10.1101/2024.09.27.614985

**Authors:** Conor P. Lane, Veronica M. Tarka, Olivier Valentin, Alexandre Lehmann, Edith Hamel, Etienne de Villers-Sidani

## Abstract

Psilocybin, a psychoactive substance derived from fungi, has been utilized historically by diverse cultures for both medicinal and non-medicinal purposes, owing to its ability to elicit profound sensory and cognitive alterations and sustain long-term changes in mood and cognition. Promising results from recent clinical studies have generated a wave of interest in employing psilocybin to treat neuropsychiatric and neuro-degenerative conditions. How psychedelics cause acute perceptual effects, and how these relate to long-lasting alterations is still debated. Whereas it is thought that perceptual disturbances may be caused by disrupted flow of information between sensory and higher order areas, *in vivo* studies have focused mostly on the latter. In particular, there has been little study of how psilocybin affects sensory representations in primary auditory cortex (A1). We used two-photon microscopy and wide field calcium imaging to examine how psilocybin affects A1 neuron response properties in the mouse. Administration of 1 mg/kg psilocybin prevented habituation of sound-evoked responses to repeated stimuli, maintaining overall responsiveness, bandwidth, and sound-level response thresholds after repeated stimulation. This was in contrast to marked habituation of responses and narrowing of tuning in controls. We observed no effect on overall distribution of best frequencies at the cortical level, suggesting psilocybin in A1 disrupts normal sensory gating, rather than tonotopic organization. This supports models of psychedelic action in which perceptual disturbances are driven by disrupted hierarchical sensory gating. With further research, influences of psychedelics on sensory representations could be harnessed to target maladaptive sensory processing in conditions such as tinnitus.

**Significance Statement:** Despite its role in altering auditory sensory perception, the impact of psilocybin on modulating neuronal activity in the auditory cortex remains understudied. This study is the first to identify an inhibition of normal auditory habituation to repeated stimuli with single-neuron resolution. We identify a role for psilocybin in the targeted, context-dependent modulation of auditory sensory neural tuning properties, which may help to explain how disruption of hierarchical control of sensory representations leads to perceptual disturbances. With further work, this influence on sensory representations could be used to target conditions where maladaptive sensory processing leads to deleterious health outcomes.

## Introduction

Serotonergic psychedelics such as psilocybin have long been employed by diverse cultures for spiritual and medicinal purposes^1^, producing profound alterations in perception and long-term changes in mood and cognition^2^. Promising results from several clinical trials have accelerated clinical interest in psychedelics to treat conditions such as depression^3–6^, addiction^7–10^ and anxiety^3,11–14^. A number of animal studies have identified a rapid induction of enhanced neuroplasticity with psychedelics, facilitating structural and functional remodeling of neural circuitry^15–19^. Despite impressive clinical findings, it remains unclear how psychedelics influence neuronal activity to exert their acute effects on sensory perception and the long-term effects that may support clinical use.

Psilocybin and its metabolite psilocin belong to the tryptamine class of psychedelics, which possess a broad affinity for a range of both serotonergic and non-serotonergic receptors throughout the brain^20^. They exhibit high affinity for both the primarily excitatory 5-hydroxytryptophan type 2A (5HT_2A_) and inhibitory 5HT_1A_ receptors, reflecting a complex pharmacological profile in which effects on neuronal excitability differ based on brain region^21,22^, neuronal subtype and dosage^21^. Indeed, psychedelics show mixed, yet primarily excitatory effects in frontal cortices^15,21,23^, and effects in sensory cortices where prior firing rate and subtype of the observed neuron determine their effect on excitability^22,24^. Multiple models of psychedelic action posit that the perception-altering effects of psychedelics are elicited through disruption of control of the flow of sensory information between sensory and higher order regions, though there is little consensus as to whether this represents over-activity or reduction of hierarchical control^20,25–27^. Neuroimaging work in human auditory cortex has found that 5HT_2A_ agonists induce acute changes in responses to sensory input, and reduce habituation to repeated stimuli^28^. In human visual cortex, they have been shown to acutely broaden visual neuron tuning^29^ and show primarily suppressive effects on excitability in mouse primary visual cortex (V1), as well as loss of surround-suppression^24^. Inhibitory regulation of sensory-evoked responses, for example by interneurons, confers context dependent-modulation of sensory neuron tuning^30–32^. These mechanisms are vital for processes such as attention, allowing for filtering of behaviorally irrelevant information^33^. These findings suggest that effects of psychedelics may involve disruption of this context-dependent modulation of sensory gating and neuronal tuning properties. This could prove useful in treatment of conditions where maladaptive plasticity of sensory representations and disruption of local cortical inhibition cause deleterious outcomes, such as tinnitus^34,35^.

The effect of psilocybin on auditory cortex sound-evoked responses and neuronal tuning properties is understudied relative to other regions, despite auditory stimulation in the form of music and chanting playing a significant role in traditional and clinical psychedelic use^36^. In particular, no study has longitudinally tracked effects on a consistent cell population at single-neuron resolution. Here, we used wide field calcium imaging and two-photon microscopy in awake, head fixed mice to assess effects of psilocybin on primary auditory cortex neuronal activity. We also tracked a subset of cells between recordings to longitudinally measure changes in individual A1 neuron tuning properties. We found that repeated stimulation in controls induced a habituation of A1 responses to sound, with a reduction in overall neuronal responsiveness, loss of responses at lower sound levels, and narrowing of tuning bandwidth. In contrast, administration of 1 mg/kg psilocybin prevented the reduction in excitatory neuronal activity and preserved baseline response thresholds and tuning width. Our findings suggest that psilocybin disrupts inhibitory habituation and filtering of familiar stimuli in A1, supporting models of psychedelic action in which perceptual disturbances are caused by loss of top-down hierarchical control over sensory gating.

## Results

### Imaging neuronal calcium responses to sound in A1

A1 was identified in wide field by the presence of a large, rostro-caudal high-to-low frequency tonotopic gradient, and from pial vessel positions (Figure 1A, B). Mice were exposed to randomly presented pure tones (4-45 kHz, 30 – 80 dB SPL) and neuronal ROIs were extracted based on calcium activity (Figure 1C) and categorized as ‘responsive’ if their activity was significantly modulated by either frequency or intensity (Figure 1D) of sound stimulation. Across all animals, 3886 neurons were identified in the pre-saline recording and 3593 in the post-saline recording. In the pre-psilocybin condition, we identified 4009 neurons, and in post, 4153. Tuning curves were reconstructed for all responsive cells (Figure 1E) and a subset of neurons was matched across recordings, in order to longitudinally examine changes in their tuning properties. We tracked 1071 neurons across the two saline session recordings, and 1831 with psilocybin. Briefly, each mouse underwent two imaging sessions separated by 5 days (N = 8 mice). In the first session, animals were exposed to an initial randomized stimulus battery (see Methods) while recording A1 activity. Immediately following this, mice were injected with 0.1 ml/10 g saline and a second recording was taken following a 15-minute wait period. The second session proceeded in the same way, with 1 mg/kg psilocybin between recordings (Figure 2A).

**Figure 1.**
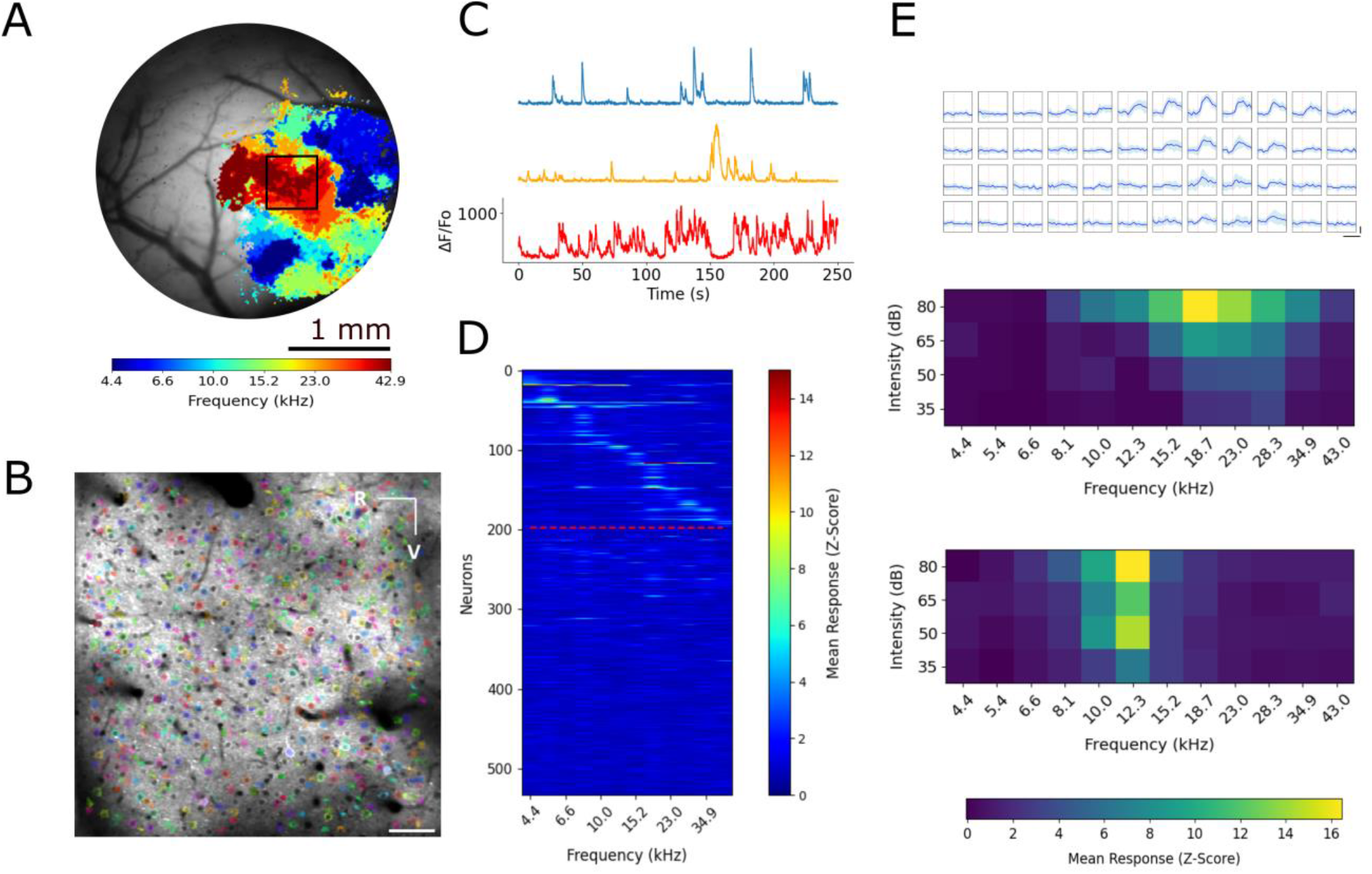
Imaging auditory cortex neuronal activity in awake mice. A) Representative best frequency tonotopic map used to locate A1, by the presence of a large, rostro-caudal high to low frequency tonotopic gradient and position of pial vessels. Inset: two-photon field of view seen in B. B) Representative frame-averaged image of mouse A1 overlaid with neuronal cell body identification. FOV is 794×794 µm, 175 µm below pial surface (layer II/III). R and V represent rostral and ventral directions. ROI coloration is random. Scale bar is 100 µm. C) Three example activity traces during sound stimulation, from ROI’s identified in B. y-axis is fractional fold-change in fluorescence from baseline (ΔF/F0), corrected for neuropil contamination. D) Trial-averaged responses of all identified neurons in B to pure tones. Cells above the red line showed activity significantly modulated by frequency or intensity (p < 0.05, two-way ANOVA), and are sorted by BF, cells below line are unsorted. E) Representative frequency-intensity tuning curves of two neurons identified in B. Above: Trial averaged (ΔF/F0) neuronal response traces. Blue line represents the mean response across trials, shading represents standard deviation. Horizontal scale bar is 1 second, vertical is a ΔF/F0 change of 100. Below: Trial-averaged Frequency-intensity tuning maps. The top map is the tuning map of raw traces shown above. Constructed from the deconvolved neuronal activity (see Methods).

**Figure 2.**
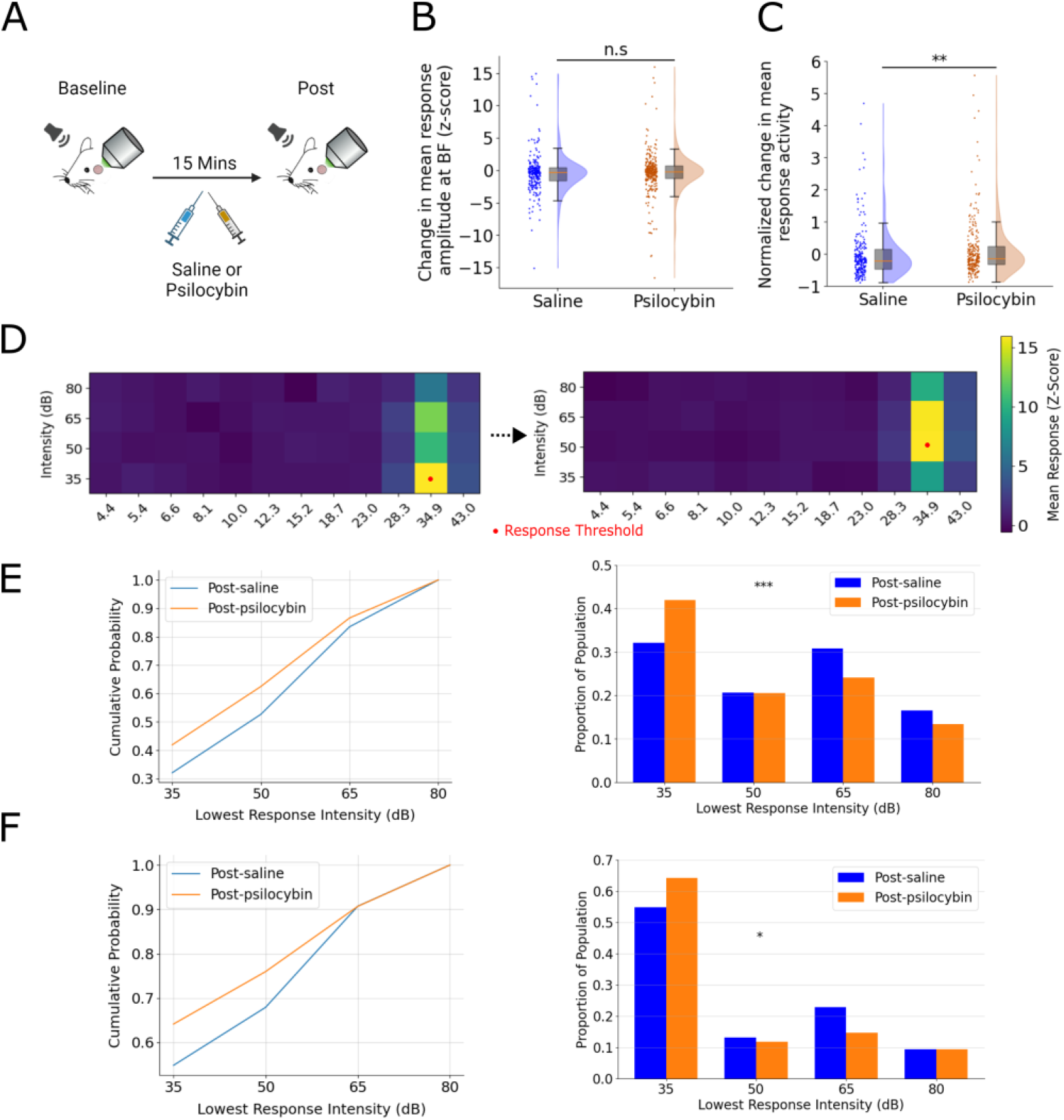
Overall response activity and sound-level response thresholds are preserved by psilocybin. A) Experimental paradigm for a single session. Each mouse has two recording days. An initial stimulus battery (see Methods) is presented, animal is then given saline IP (session 1) or 1 mg/kg psilocybin (session 2) and a second recording taken following 15 min incubation. 5 days are left between saline and psilocybin recording sessions and the same cell population is recorded from across all four recordings. B) Change in trial-averaged peak response amplitude at BF, for matched, consistently responsive cells. n.s, p = 0.120, (Mann Whitney U). C) Change in mean firing activity (deconvolved activity, see Methods) across all frequency intensity combinations, between pre- and post-recordings, for matched, consistently responsive cells. Relative change is normalized to baseline response values for each neuron. Shaded area represents kernel density estimate of distribution. Boxplot, orange line is median, grey shading is interquartile range, whiskers are SEM. **, p = 0.006, Mann Whitney U. D) Representative cell tuning matched across recordings, illustrating a rise in lowest response threshold with saline. E) Left: Cumulative distribution plots of the lowest sound intensity eliciting an above threshold response, for all responsive neurons in the population. Distributions show the post-drug intervention recording for saline and psilocybin conditions. Right: Distribution of lowest response threshold for post-saline and post-psilocybin conditions, for all responsive neurons in population. Normalized according to total cell number in each group. ***, p = 4.12×10^−5^ (Mann Whitney U). N Post saline = 1222 cells, post psilocybin = 1194 cells. F) Left: lowest sound intensity, for cells matched and consistently responsive across pre- and post-recordings. Right: Distribution of lowest response intensity for matched, consistently responsive cells. *, Mann Whitney U, p = 0.040. B, C, F) N post saline = 237 cells, post-psilocybin = 328.

### Sensitivity to low sound levels following repeated stimulation is preserved with psilocybin

In consistently responsive, matched cells, we observed a significant difference in the change in median trial-averaged response amplitude, averaged across frequency-intensity combinations, between saline and psilocybin conditions (p = 0.004, Mann Whitney U). Saline showed a median reduction of -0.211 and psilocybin, -0.133 (z-score) (Figure 2C). However, when examining the trial-averaged mean response amplitude at best frequency (BF), we observed no difference between saline and psilocybin conditions (Figure 2B) (p = 0.133), despite saline showing a trend change of -0.327 and psilocybin, -0.148. This suggested that reduction in overall response activity in saline may not be due primarily to a change in the amplitude of responses at BF, but instead may be due to shifts in the off-BF shape of tuning curves. To investigate this further, we examined changes in sound-level response thresholds of neurons with repeated stimulation (Figure 2D). Across all responsive cells (Figure 2E, left) and in the matched cell subset (Figure 2F, left), those in the post-psilocybin condition exhibited a higher cumulative probability of responding to lower intensity stimuli, indicating a preservation of sound-level response thresholds. We observed a significant difference in the distribution of response thresholds between the post-saline and post-psilocybin recording, both in all active cells (p = 4.12 x10^−5^, Mann Whitney U) (Figure 2E, right) and in matched cells (Figure 2F, right) (p = 0.04). No significant difference in lowest response intensity was observed between the pre-drug recordings for saline and psilocybin conditions in all (p = 0.12 Mann Whitney U), or matched cells (p = 0.86) (Figure Supplementary 2A, B respectively), ensuring there was no confounding effect of performing both recording sessions in the same animal. These findings suggest that administration of 1 mg/kg psilocybin prevents the habituation in overall responsiveness and raising of response thresholds that occurs with repeated exposure to the stimulus.

### Narrowing of tuning bandwidths with repeated stimulation is inhibited by psilocybin

We measured changes in the bandwidth tuning (Figure 3A) of neurons at a range of sound intensities. At 80 dB, we observed no significant difference in the distribution of response bandwidths pre- and post-intervention in either the saline (p = 0.671, Mann Whitney U) or psilocybin (p = 0.288) conditions, although data suggest a trend towards a reduction in the saline condition (Figure 3B). At 65 dB, however we observed a significant reduction in response bandwidths in the saline condition (p = 0.006) that was not present in the psilocybin condition (p = 0.947) (Figure 3C). A similar reduction was also observed at 50 dB with saline (p = 0.015) but not with psilocybin (p = 0.199) (Figure 3D). Finally, no difference was found at 35 dB between the saline (p = 0.300) and psilocybin (p = 0.596) conditions, although a trend towards a reduction was observed with saline (Figure 3E). At all intensities, no significant difference was observed in the pre-drug distributions of bandwidths between saline and psilocybin conditions (data not shown, p > 0.190 for all). These data suggest that the narrowing of tuning occurring with repeated stimulation is prevented by psilocybin administration, maintaining a state in A1 as if responses are to a novel and not a familiar stimulus.

**Figure 3.**
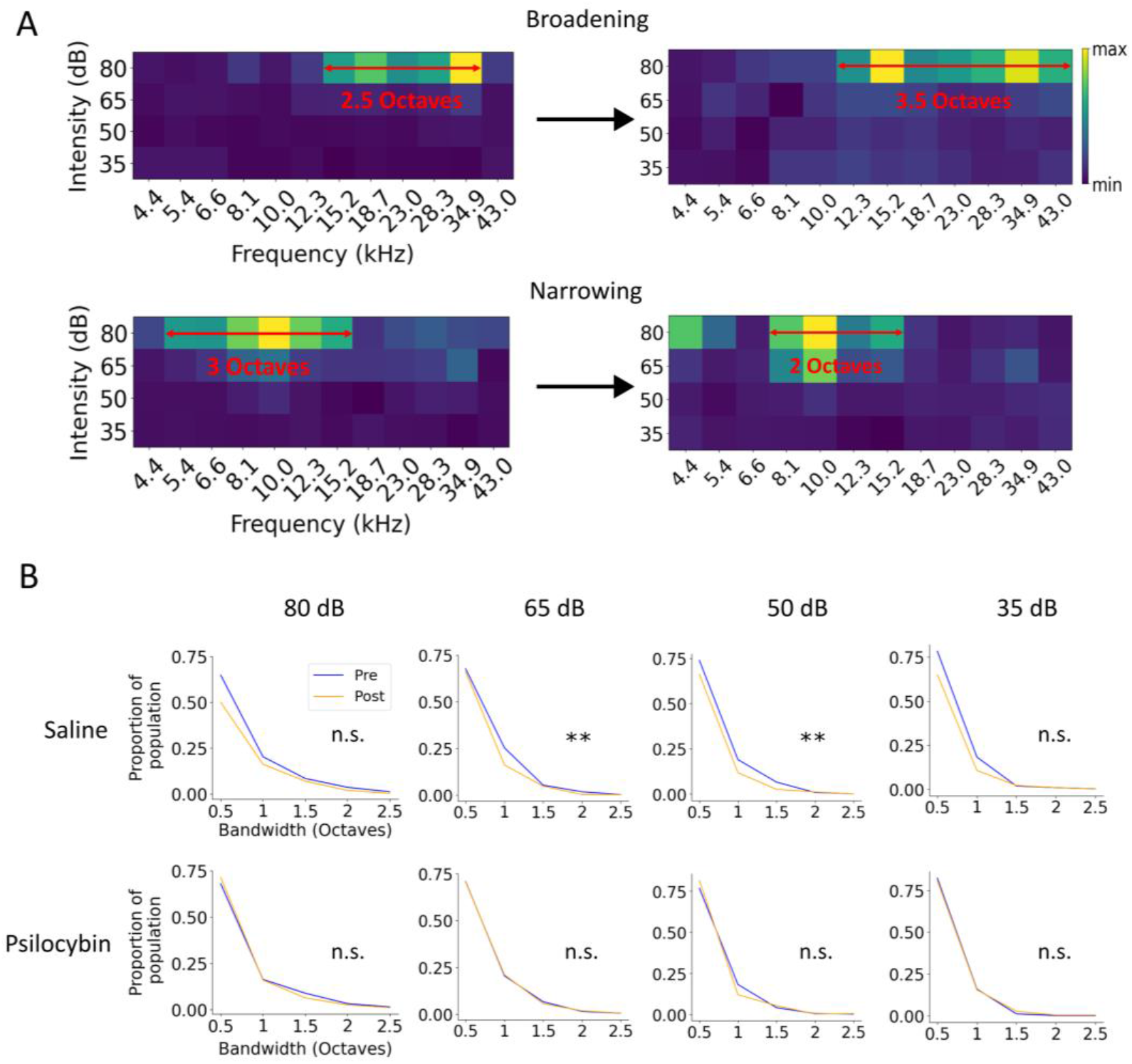
Narrowing of tuning bandwidths with habituation is prevented by psilocybin. A) Representative tuning curves from matched cells pre- and post-drug intervention, illustrating a broadening of 80 dB bandwidth (above) or narrowing (below) in a single cell between recordings. Bandwidth is the continuous range of frequencies at the given intensity, at which an at least half-maximal response is observed. B) Change in bandwidth at 80, 65, 50 and 35 dB SPL between pre- and post-drug intervention recordings, for saline condition (above) and psilocybin condition (below). Data are normalized as relative frequency of each bandwidth occurring within that group. n.s = p > 0.05, ** = p < 0.01. Saline left to right: N pre-saline = 540, 536, 401, 317 cells, respectively. Post = 412, 467, 326, 251 cells. Psilocybin left to right: N Psilocybin pre = 391, 425, 334, 283 cells. Post = 343, 394, 332, 304.

### Rapid drift in neuronal response properties is observed, and is unaffected by psilocybin

Underlying relatively stable tonotopic representations in macro-cortical space, individual neuron tuning properties shift constantly, a phenomenon called representational drift^37^. We hypothesized that psilocybin’s acute effects on neuronal activity may alter the rate of this drift. Of tracked neurons, a sizeable proportion shifted in or out of sound-sensitivity between recordings (Figure 4A), with fewer than half responsive in both. No significant differences between saline or psilocybin conditions were observed across any of the response categories (Mann Whitney U, Supplementary Figure 2). We observed a pronounced shift in preferred intensity (PI) in both the saline and psilocybin recordings (Figure 4B), with 35.9% of cells shifting PI by 10 dB or more in the saline condition, and 38.7% with psilocybin. For the distributions of both the absolute magnitude of the PI shifts (data not shown) and signed change, no difference was observed between saline and psilocybin (p = 0.446, p = 0.700, respectively, Mann Whitney U). Similarly, a sizeable proportion of cells showed a shift in BF between recordings (Figure 4C), with 48.9% shifting by 0.5 octaves or more with saline, and 45.9% with psilocybin. No difference was observed between saline and psilocybin sessions (p = 0.374, p = 0.110 respectively, Mann Whitney U) for magnitude or signed change. To ensure that observed changes were not due to degradation in the signal-to-noise ratio across recordings, we computed the trial-trial reliability (Quality Index, see Methods). We found no difference in the matched-cell shift in reliability between pre- and post-recordings for saline and psilocybin groups (p = 0.120, Kolmogorov-Smirnov) (Figure 4D, E). Taking the whole cell population, we observed no change in the distribution of response reliabilities in either the saline or psilocybin conditions (p = 0.414, p = 0.339, respectively, Kolmogorov-Smirnov) (Figure Supplementary 2C, D). We also examined the distribution of BFs at the whole-population level (Figure 4F), averaged across mice (N = 8). We identified no significant changes in BF’s between pre- and post-recordings, for either the saline or psilocybin condition (Mann Whitney U and Bonferroni). Taken together, these data suggest a significant and rapid degree of representational drift in single-cell tuning properties over a notably short period of time, while maintaining stable representation at the cortical level. The magnitude of drift appears to be unaffected by psilocybin.

**Figure 4.**
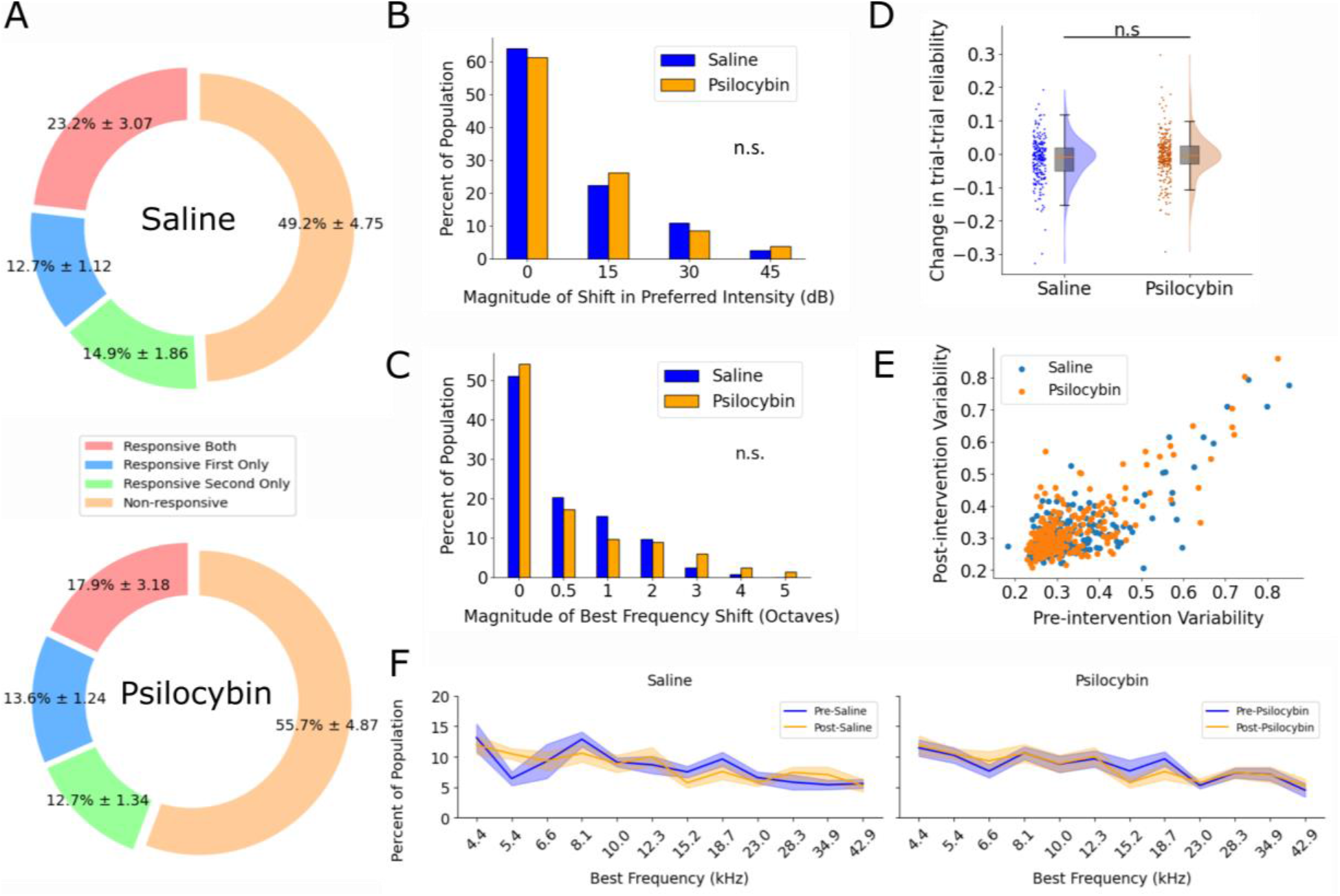
Rapid drift in neuronal tuning properties is observed and is unaffected by psilocybin. A) Categorization of when neurons were sound-sensitive as a percentage of the overall population (% ±, SEM), when tracked across recordings. Cells were categorized as either responsive to sound in both recordings, responsive only in the first recording, responsive only in the second, or non-responsive in either recording. No significant differences in proportion between saline and psilocybin conditions were observed (Mann Whitney U). N = 8 mice. B) Change in preferred intensity (intensity at which the maximum trial-averaged response was elicited) across cells responsive in both recordings. Difference between the two distributions is non-significant (Mann Whitney U, p = 0.70). C) Magnitude of shift in neuronal best frequency between recordings, across matched cells responsive in both recordings. Difference between the two distributions is non-significant (Mann Whitney U, p = 0.10). B, C) N Saline = 237, N Psilocybin = 328 cells from 8 mice. D) Shift in trial-trial reliability for matched, consistently responsive cells. n.s, p = 0.120, Mann Whitney U. Shaded area represents kernel density estimate of distribution. Boxplot, orange line is median, grey shading is interquartile range, whiskers are SEM. E) Distribution of pre- and post-intervention trial-trial reliability for matched, consistently responsive cells. A-E) N Saline = 237, N psilocybin = 328 cells from 8 mice. F) Distribution of BFs in the neuronal population, as a percentage of total responsive cells (unmatched). Shaded areas are mean ± SEM. No significant difference was observed between pre- and post-recordings for any BF, in either saline or psilocybin conditions. No significant difference was observed in the overall distribution in the post-recording, between saline or psilocybin (Mann Whitney U and Bonferroni multiple comparisons correction). N pre-saline = 1307 cells, N post = 1222. N pre-psilocybin = 1194 cells, N post = 1214.

## Discussion

This study identified a number of key findings: 1) Psilocybin prevents habituation of neuronal response activity 2) Psilocybin preserves neuronal response thresholds and prevents narrowing of frequency tuning with habituation, while maintaining population-level BF tuning. 3) Rapid representational drift occurs in A1 on the order of minutes, which is unaffected by psilocybin. Our results indicate a clear effect of psilocybin on sensory gating in A1.

### Psilocybin prevents habituation of neural firing activity to repeated stimulation

We observed a marked habituation of responses across the two control recordings, in terms of overall neuronal firing activity, narrowing of tuning bandwidth and loss of sensitivity to lower intensity sounds (Figure 2C, E, F. 3C, D). Similar effects on responses with repeated stimulation have been observed in awake mouse A1^30,38^ and in V1^39^, remaining under-reported until recently as the effect is neutralized by anesthesia^30^. The fact that amplitude of responses at BF was unaffected (Figure 2B) suggests that reduction in response activity came from loss of responses to off-BF frequencies and off-PI sound-levels. In contrast, we observed no reduction in firing activity with psilocybin, indicating that a state where the stimuli was perceived as novel was preserved by the drug. A recent two-photon study^40^ reported a decrease in neuronal responsiveness in A1 30 minutes following psilocybin, though their data show no significant reduction from baseline at this time-point. As our recordings cover the 30-minute time-point, we suggest this does not contradict our finding. A recent study using 2,5-Dimethoxy-4-iodoamphetamine (DOI) in V1 reported a net-suppression of activity, with suppression or facilitation depending on prior neuronal firing rate^24^. We did not observe suppression, which may reflect differential effects of sensory gating between cortices, for example opposing effects of locomotion on evoked activity in auditory^41^ versus visual cortex^42^. It may also be due to drugs used, as psilocybin and DOI belong to the tryptamine and phenethylamine classes, respectively^43^, with different receptor affinities. Our early findings therefore highlight the interesting prospect that rather than shifting response gain, psilocybin may affect modulation of tuning curve width and sound-level selectivity as stimuli become familiar.

### Psilocybin prevents rise in sound-level response thresholds and narrowing of tuning with repeated stimulation

We observed a significant difference in the distributions of intensity-response thresholds between control and psilocybin conditions, with psilocybin exhibiting greater maintenance of responses to low sound intensities. This effect was observed both at the population level (Figure 2E), and in consistently responsive, tracked cells (Figure 2F). No difference between control and psilocybin groups was observed for response thresholds before intervention (Figure Supplementary 2A-B), suggesting that at least in layer II/III, habituation returned to baseline after five days, and that performing saline controls in the same animal did not confound psilocybin-session results. The difference was most pronounced at lower sound intensities (35, 50dB), which may be explained by increased adaptation previously observed at lower sound intensities in auditory cortex^44^ and in inferior colliculus^45^. These data indicate that psilocybin prevents the loss of sensitivity to low sound levels following repeated stimulation. Habituation of responses during repeated stimulation also narrows frequency tuning of neurons in primary auditory cortex^46^, and studies have highlighted the role of increasing inhibition in sharpening tuning^32,47^. We observed this narrowing with repeated stimulation as significant at the mid-range of presented sound levels, namely 65 and 50 dB (Figure 3C, D) but not at the lowest and highest levels, 35 and 80 dB (Figure 3B, E), although a trend towards narrowing of tuning was present. Importantly, this was not observed at any sound level following psilocybin administration, suggesting that it prevents inhibition-driven narrowing of tuning.

### Rapid representational drift is observed in A1, which is unaffected by psilocybin

We found a high degree of drift in BF and PI between pre- and post-recordings in both control and psilocybin conditions, but the rate of drift was unaffected by psilocybin (Figure 4 B-C). In both conditions, less than half of sound-responsive neurons were so across both recordings (Figure 4A), again with no differences between control or psilocybin conditions (Supplementary Figure 1). The scale of representational drift observed is similar to previous studies in A1^48^, but the rate is of note. Where it was previously measured on the order of days, our data show that this instability is pronounced on the order of minutes, in line with findings in the visual cortex^49^. We observed no change in population-level BF tuning with psilocybin (Figure 4F), supporting previous findings that macro-cortical preferred representations are preserved^24,40^, suggesting that the effects of psilocybin involve disruption of sensory gating, rather than overall tonotopic organization.

## Conclusions

By preventing habituation in primary auditory cortex, our findings highlight a much discussed aspect of psychedelic action, the relaxation of top-down predictive control. This is supported by findings with other psychedelics and in other sensory modalities. A magnetoencephalography study in humans found that LSD reduces response adaptation in A1^28^. Studies using DOI^24^ and DMT^29^ also showed loss of surround suppression and broadening of visual receptive fields. In sensory cortices, stimulus novelty and behavioral relevance are encoded through inhibitory, interneuron-driven modulation of sensory neuron tuning^30,32^. In particular, somatostatin interneurons regulate cortical habituation^30,31^, and surround suppression in V1^50^. Given that GABAergic interneurons express 5HT_2A_ receptors^51^ and their activity is modulated by tryptamine psychedelics^52,53^, it is interesting to speculate that this loss of habituation may be due to direct action of psilocybin on cortical interneurons. A ‘synthetic surprise’ model of psychedelic action recently proposed by De Filippo and Schmitz^26^ argues that perceptual alterations may be caused by influencing interneuron-driven encoding of prediction error^54^. Future studies should examine more directly the effect of psilocybin on interneuron activity in A1. Moreover, direct examination of psilocybin’s effect on familiar versus unexpected stimulus responses (e.g. auditory oddball paradigm^55^) could confirm whether prediction-error responses are disrupted.

Our research in A1 reveals significant insights into the neurophysiological actions of psilocybin. By preserving auditory response thresholds and inhibiting the habitual narrowing of neuronal tuning, our study underscores the influence of psilocybin on sensory gating and neural plasticity. Importantly, our study tracked neurons longitudinally across recordings, highlighting that the effect of psilocybin is present at the level of individual cells and at the population-level. By maintaining heightened sensory responsiveness, psilocybin may promote a state of “neural openness,” which facilitates the reconfiguration of perceptual and cognitive hierarchies critical for adaptive behavior and mental health. This aligns with theoretical frameworks suggesting that the therapeutic effects of psychedelics stem from their capacity to foster a flexible mental state^25^. The ability of psychedelics to disrupt entrenched sensory and cognitive processing holds significant therapeutic potential, especially in treating conditions characterized by rigid, maladaptive neural circuits such as in depression, addiction, and PTSD. A deeper understanding of these mechanisms could pave the way for using psychedelics in a controlled and therapeutically beneficial manner, offering new hope for numerous patients whose conditions are resistant to conventional treatments.

## Materials and Methods

### Animals and Drugs

All experimental procedures were approved by the Montreal Neurological Institute Animal Care Committee and follow guidelines of the Canadian Council on Animal Care. Experiments were performed on male and female adult mice (no older than 24 weeks) of the Thy1-GCaMP6s strain (Jackson labs, stock number 024275), crossed with CBA/CaJ (Stock No. 000654), a strain that retains good hearing thresholds^56^. GCaMP6s-positive offspring retain good hearing thresholds well into adulthood^57^. Mice were maintained on a 12-hour light/dark cycle with *ad libitum* access to food and water and were group-housed until cranial window surgery. Psilocybin was supplied by Psygen Inc. (Calgary, AB) and use of psilocybin was approved by Health Canada, April 2022.

### Cranial Window Surgery

Cranial window procedure was adapted from Romero *et al 2020*^57^. Briefly, animals were anaesthetized with isoflurane in oxygen (5% induction, 1-2% maintenance). An incision was made to expose the skull and periosteum removed from the skull. Custom head-fixation hardware (iMaterialise, Belgium) was attached to the skull using dental cement (C&B Metabond). Left auditory cortex was localized using skull anatomical landmarks and a 3 mm craniotomy drilled. A 3 mm glass coverslip (#0 thickness, Warner Instruments, MA) was then inserted into the craniotomy and affixed with dental cement. Animals were allowed to recover in a warmed chamber and administered 20 mg/kg carprofen for three days post-operatively. Imaging began a minimum of 5 days following surgery.

### Two-photon Imaging

Imaging of calcium activity was performed using an Ultima Investigator two-photon microscope (Bruker, MA) with a Cambridge Technology 8 kHz CRS resonant scanner (Novanta Photonics, MA) and 350-80LA pockels cell (Conoptics, CT). Excitation light was provided using a Mai-Tai eHP laser (SpectraPhysics, UK) tuned to 920 nm. System was controlled and images collected with PrairieView software (Version 5.4, Bruker). Microscope and animal fixation set-up were enclosed in a light-attenuating box. Resonant scanner was used at a resolution of 512×512 pixels, collecting images at 10 Hz. A 16X/0.80 LWD immersion objective (Nikon, UK) was employed. Neuronal activity was imaged at 175 μm below the pial surface (layer II/III).

### Wide field Imaging

Wide field calcium activity was collected using a PCO Panda 4.2 scMOS camera (Excelitas Technologies, MA) at 10 Hz framerate. A UPlanFL N 4X, 0.13 numerical aperture lens (Olympus, Japan) was used. Excitation light was provided by an X-Cite Series 120 Q lamp (Excelitas Technologies) and passed through a 470-10 nm band pass filter (Thorlabs, NJ). Light was focused at a depth of 200 μm. Images were collected using PCO Camware software (version 4.12, Excelitas).

### Imaging procedure

Prior to imaging, animals were acclimatized to head fixation for three days (15, 30 and 45 minutes). On imaging day 1, a wide field tonotopic map was taken to ascertain the location of primary auditory cortex (A1) and the two-photon field of view fixed here. Both saline and psilocybin sessions were conducted in the same animal, so that animals underwent two imaging sessions separated by 5 days. On each day, batteries of stimuli were presented, immediately followed by intraperitoneal (IP) injection of saline or 1 mg/kg psilocybin (equal volumes) and a 15-minute wait period. To avoid potential long-term effects of psilocybin, all animals were administered saline on session 1 and psilocybin on day 2.

### Sound presentation

Stimuli were designed with OpenEx software (Tucker-Davis Technologies (TDT), FL) and generated using a TDT RZ6 multi I/O processor (TDT). Sounds were presented with an MF1 free-field speaker (TDT) 10 cm from the contralateral (right) ear. ***Two-photon:*** Neuronal receptive fields were reconstructed using responses to a range of frequency-intensity combinations of pure tones. For two-photon imaging, 12 sound frequencies (100 ms tones, 5 ms on/off ramps, ranging from 4-45 kHz in 0.5 octave increments) were presented at four intensities (35-80 dB SPL, 15 dB increments), with a 1.5s inter-stimulus interval (ISI). Stimuli were presented in random order, with each of the 48 total frequency-intensity conditions repeating 10 times. ***Wide field:*** To reconstruct wide field tonotopic maps, 12 frequencies were presented at 65 dB SPL (100 ms tones, 5 ms on/off ramps, ranging from 4-45 kHz in 0.5 octave increments), each repeating ten times with a 2.5s ISI.

### Data analysis – Two-photon

#### Extracting neuronal calcium responses from two-photon recordings

Recordings were imported into Suite2p^58^, an open source package providing a pipeline for extraction of calcium signals from neuronal ROI’s. Suite2p performs motion correction, extraction of calcium traces and deconvolution to infer spike activity. Detected neuronal ROI’s were manually confirmed as healthy neurons by the presence of a clearly demarcated cell body, a ring of fluorescence and a dark center. The neuronal calcium trace (ΔF/F0) and deconvolved spike activity were then exported for further analysis in Python.

#### Identification of sound-sensitive neurons

The total trial length was 1.5s (−0.5 seconds from stimulus onset to 1s post-onset) and response period was stimulus onset to 1s post-onset. Evoked activity was defined as the median trial-averaged activity during the response period. We determined whether a cell was sound-sensitive using two-way ANOVA, with tone frequency and sound intensity of as predictors. Cells significantly modulated by frequency or intensity (p < 0.05 main effect for either frequency, or frequency/intensity interaction) were labelled as sound-sensitive and taken forward for further analysis.

#### Matching neurons across recordings

In order to directly examine changes of tuning with single-cell resolution, a subset of identified neurons was tracked across the pre- and post-recordings. ROI coordinates and mean image for each recording were imported to the open source MATLAB package ROIMatchPub, which displays aligned ROI’s for manual confirmation.

#### Calculating individual neuron tuning properties

Frequency-intensity tuning curves were reconstructed from the peak of the trial-averaged baseline-corrected response. *Best frequency (BF), Preferred intensity (PI):* BF and PI were calculated as the frequency or intensity of stimulation eliciting the maximal trial-averaged peak response, respectively. *Bandwidth:* Bandwidth was defined as the continuous range of frequencies around the BF with a response above half-maximum. *Lowest Response Intensity:* The lowest response intensity was defined as the lowest intensity at which an above threshold (4 SD from baseline) peak response was elicited. *Trial-Trial Variability:* We used a response quality index (QI), described in detail in^59^ to compute signal-to-noise ratio:

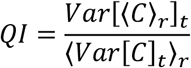

where *C* is the *T* by *R* response matrix (time samples by stimulus repetitions) and ⟨ ⟩x and Var[]x denote the mean and variance across the indicated dimension, respectively. A higher QI denotes that the mean response is a closer representation of variance across trials, with a QI of 1 being a perfect representation.

### Data Analysis – wide field

#### Image pre-processing

Recordings were motion corrected by aligning all frames to a mean reference image, constructed from the 20 most correlated of a random selection of frames. Recordings were then down-sampled from 1024×1024 pixels to 256×256. Global fluctuations in the signal were removed by fitting a linear regression model, which predicts pixel intensities of each frame based on global average intensity, and subtracts predicted global fluctuations from real activity. Trial length was 2.5s (−0.5s from stimulus onset to 2s post-onset). Pixel traces were baseline corrected by subtracting the mean of the five pre-stimulus baseline frames from the mean of the response period, which was taken as the 0.8s following stimulus onset. Pixel responses were then expressed as z-scores from pre-stimulus baseline.

#### Tonotopic map reconstruction

Best Frequency (BF) tonotopic maps were reconstructed first by z-scoring median pixel responses to a given frequency relative to every other pixel. This corrects for biases in response amplitude in the cortex for particular frequency ranges. Only responses with a median z-score greater than 1 were included in analyses. The frequency at which the maximal trial-averaged median response was elicited was defined as the BF.

### Quantification and statistical analysis

Analyses were conducted with Python 3.12. Data are reported as mean ± SEM unless stated otherwise. For categorization of cell sound-sensitivity, two-way ANOVA was used. For comparison between groups, Mann Whitney U or Kolmogorov-Smirnov tests were used. Statistical significance was defined as p < 0.05.

## Acknowledgments

We thank Dr. Daniel B. Polley for sharing head holder 3D print files, and Dr’s Anne E. Takesian and Maryse E. Thomas for supporting establishing cranial window protocol. Thank you to Dr. Stuart Trenholm for support establishing two-photon analysis pipelines. Psilocybin was supplied by Psygen Inc., Alberta. The following funding sources supported this study: Fonds de recherche du Québec - Santé, Doctoral Scholarship. Centre for Research on Brain, Language and Music, Research Incubator Award to A.L and E.V.S. Canada Institutes of Health Research Project Grants to E.V.S and E.H.

## Supplementary Information

**Supplementary Figure 1.**
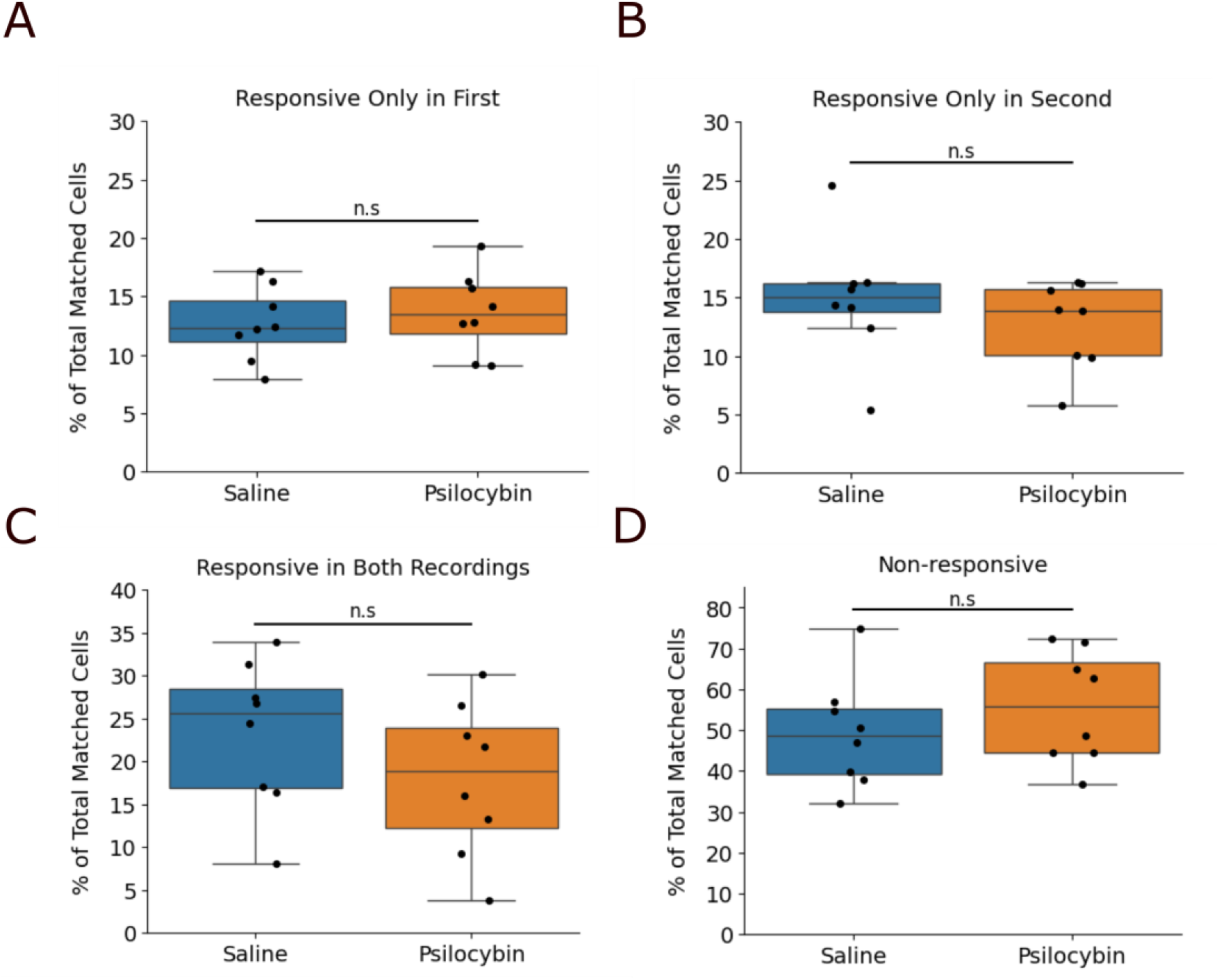
No difference in whether cells were sound-sensitive across recordings, between saline and psilocybin conditions. Response characterization of cells tracked across pre- and post-intervention recordings, as a percentage of the total number of tracked cells in that recording. A) Cells responsive to changes in frequency/intensity only in the first recording. B) Cells responsive only in the second recording. C) Cells consistently responsive in both recordings. D) Cells responsive in neither of the two recordings. n.s = p > 0.05, Mann Whitney U test. Boxplot, middle line is median, colored shading is interquartile range, whiskers are SEM.

**Supplementary Figure 2.**
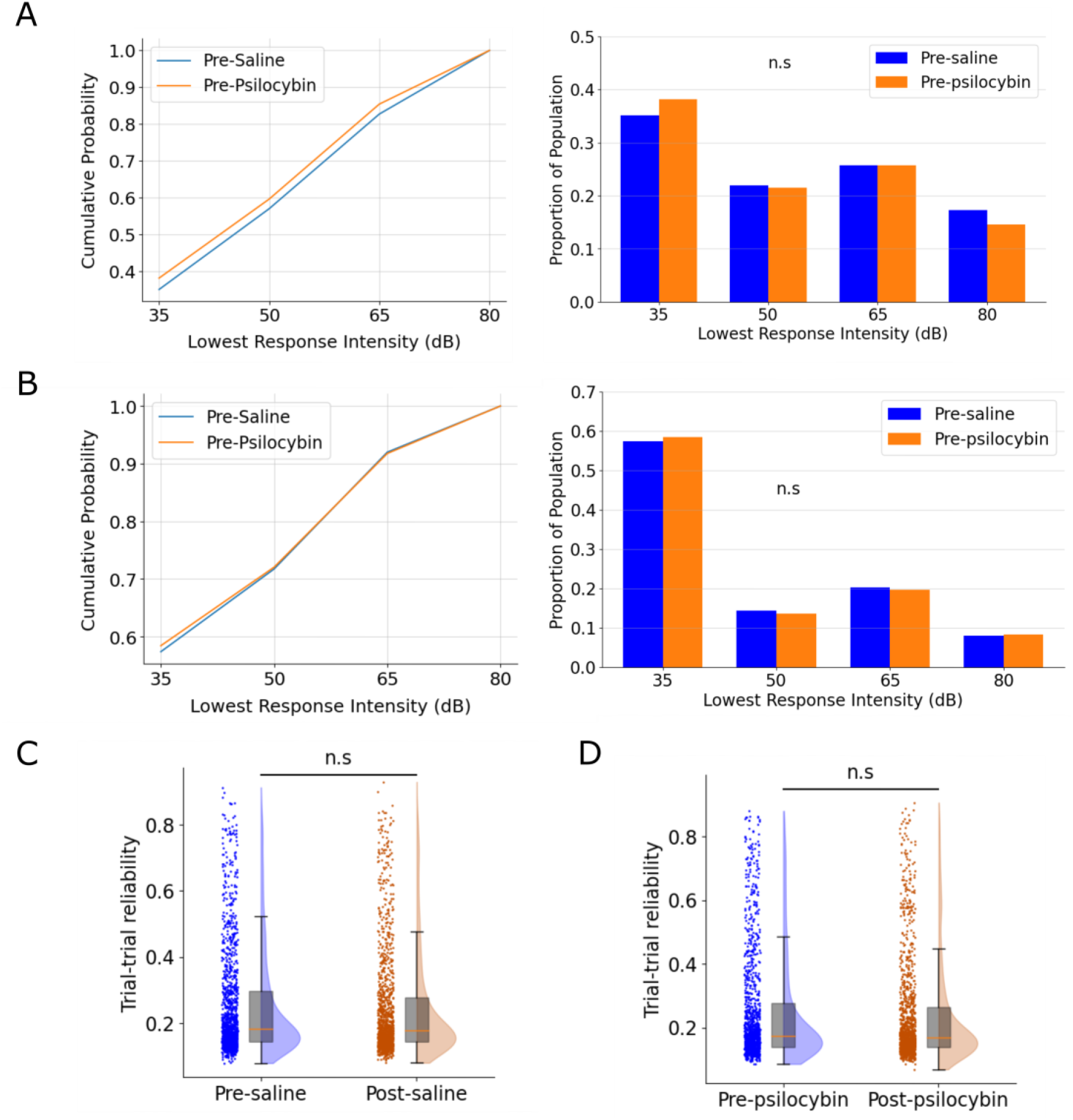
No difference in baseline response threshold and reliability between saline and psilocybin conditions. A) Left: Cumulative distribution comparing the pre-drug intervention lowest response threshold values for the saline and psilocybin conditions, for all responsive cells in the population. Right: Distribution of lowest response threshold values. B) Left: Cumulative distribution comparing the baseline lowest response threshold values for the saline and psilocybin conditions, for matched cells responsive in both recordings. Right: Distribution of lowest response threshold values C) Trial-trial reliability across all responsive cells, for the pre- and post-saline conditions. D) Trial-trial reliability for cells tracked across recordings, for the pre- and post-psilocybin conditions. n.s. = p > 0.05, Mann Whitney U.

## References

1. Nichols, D. E. Psilocybin: from ancient magic to modern medicine. J. Antibiot. (Tokyo). 73, 679–686 (2020).

2. Erritzoe, D. et al. Effects of psilocybin therapy on personality structure. Acta Psychiatr. Scand. 138, 368–378 (2018).

3. Griffiths, R. R. et al. Psilocybin produces substantial and sustained decreases in depression and anxiety in patients with life-threatening cancer: A randomized double-blind trial. J. Psychopharmacol. 30, 1181–1197 (2016).

4. Raison, C. L., Sanacora, G. & Woolley, J. Single-Dose Psilocybin Treatment for Major Depressive Disorder: A Randomized Clinical Trial. JAMA 5;330, 843--853 (2023).

5. von Rotz, R. et al. Single-dose psilocybin-assisted therapy in major depressive disorder: A placebo-controlled, double-blind, randomised clinical trial. eClinicalMedicine 56, 101809 (2023).

6. Galvão-Coelho, N. L. et al. Classic serotonergic psychedelics for mood and depressive symptoms: a meta-analysis of mood disorder patients and healthy participants. Psychopharmacology (Berl). 238, 341–354 (2021).

7. Johnson, M. W., Garcia-romeu, A., Cosimano, M. P. & Griffiths, R. R. Pilot Study of the 5-HT2AR Agonist Psilocybin in the Treatment of Tobacco Addiction. J Psychopharmacol. 28, 983–992 (2014).

8. Johnson, M. W., Garcia-Romeu, A. & Griffiths, R. R. Long-term follow-up of psilocybin-facilitated smoking cessation. Am. J. Drug Alcohol Abuse 43, 55–60 (2017).

9. Bogenschutz, M. P. et al. Psilocybin-assisted treatment for alcohol dependence: A proof-of-concept study. J. Psychopharmacol. 29, 289–299 (2015).

10. Bogenschutz, M. P. et al. Percentage of Heavy Drinking Days Following Psilocybin-Assisted Psychotherapy vs Placebo in the Treatment of Adult Patients with Alcohol Use Disorder: A Randomized Clinical Trial. JAMA Psychiatry 79, 953–962 (2022).

11. Ross, S. et al. Rapid and sustained symptom reduction following psilocybin treatment for anxiety and depression in patients with life-threatening cancer: A randomized controlled trial. J. Psychopharmacol. 30, 1165–1180 (2016).

12. Gasser, P. et al. Safety and efficacy of lysergic acid diethylamide-assisted psychotherapy for anxiety associated with life-threatening diseases. J. Nerv. Ment. Dis. 202, 513–520 (2014).

13. Palhano-Fontes, F. et al. Rapid antidepressant effects of the psychedelic ayahuasca in treatment-resistant depression: A randomized placebo-controlled trial. Psychol. Med. 49, 655–663 (2019).

14. Sanches, R. F. et al. Antidepressant effects of a single dose of ayahuasca in patients with recurrent depression a SPECT study. J. Clin. Psychopharmacol. 36, 77–81 (2016).

15. Shao, L.-X. et al. Psilocybin induces rapid and persistent growth of dendritic spines in frontal cortex in vivo. 109, 2535–2544 (2022).

16. Nardou, R. et al. Psychedelics reopen the social reward learning critical period. Nature 618, 790–798 (2023).

17. Moliner, R. et al. Psychedelics promote plasticity by directly binding to BDNF receptor TrkB. Nat. Neurosci. 26, 1032–1041 (2023).

18. Jefferson, S. J. et al. 5-MeO-DMT modifies innate behaviors and promotes structural neural plasticity in mice. Neuropsychopharmacology 48, 1257–1266 (2023).

19. Ly, C. et al. Psychedelics Promote Structural and Functional Neural Plasticity. Cell Rep. 23, 3170–3182 (2018).

20. Kwan, A. C., Olson, D. E., Preller, K. H. & Roth, B. L. The neural basis of psychedelic action. Nat. Neurosci. 25, 1407–1419 (2022).

21. Wood, J., Kim, Y. & Moghaddam, B. Disruption of prefrontal cortex large scale neuronal activity by different classes of psychotomimetic drugs. J. Neurosci. 32, 3022–3031 (2012).

22. Riga, M. S., Bortolozzi, A., Campa, L., Artigas, F. & Celada, P. The serotonergic hallucinogen 5-methoxy-N,N-dimethyltryptamine disrupts cortical activity in a regionally-selective manner via 5-HT1A and 5-HT2A receptors. Neuropharmacology 101, 370–378 (2016).

23. Riga, M. S., Soria, G., Tudela, R., Artigas, F. & Celada, P. The natural hallucinogen 5-MeO-DMT, component of Ayahuasca, disrupts cortical function in rats: Reversal by antipsychotic drugs. Int. J. Neuropsychopharmacol. 17, 1269–1282 (2014).

24. Michaiel, A. M., Parker, P. R. L. & Niell, C. M. A Hallucinogenic Serotonin-2A Receptor Agonist Reduces Visual Response Gain and Alters Temporal Dynamics in Mouse V1. Cell Rep. 16, 4375–3483 (2019).

25. Carhart-Harris, R. L. & Friston, K. J. REBUS and the anarchic brain: Toward a unified model of the brain action of psychedelics. Pharmacol. Rev. 71, 316–344 (2019).

26. De Filippo, R. & Schmitz, D. Synthetic surprise as the foundation of the psychedelic experience. Neurosci. Biobehav. Rev. 157, 105538 (2024).

27. Doss, M. K. et al. Models of psychedelic drug action: Modulation of cortical-subcortical circuits. Brain 145, 441–456 (2022).

28. Timmermann, C. et al. LSD modulates effective connectivity and neural adaptation mechanisms in an auditory oddball paradigm. Neuropharmacology 142, 251–262 (2018).

29. Pais, M. et al. Rapid effects of tryptamine psychedelics on perceptual distortions and early visual cortical population receptive fields. Neuroimage 297, (2024).

30. Kato, H. K., Gillet, S. N. & Isaacson, J. S. Flexible Sensory Representations in Auditory Cortex Driven by Behavioral Relevance. Neuron 88, 1027–1039 (2015).

31. Kato, H. K., Asinof, S. K. & Isaacson, J. S. Network-Level Control of Frequency Tuning in Auditory Cortex. Neuron 95, 412-423.e4 (2017).

32. Isaacson, J. S. & Scanziani, M. How inhibition shapes cortical activity. Neuron 72, 231–243 (2011).

33. van Moorselaar, D. & Slagter, H. A. Inhibition in selective attention. Ann. N. Y. Acad. Sci. 1464, 204–221 (2020).

34. Shore, S. E., Roberts, L. E. & Langguth, B. Maladaptive plasticity in tinnitus-triggers, mechanisms and treatment. Nat. Rev. Neurol. 12, 150–160 (2016).

35. Møller, A. R. The role of neural plasticity in tinnitus. Textb. Tinnitus 166, 99–102 (2011).

36. Jerotic, K., Vuust, P. & Kringelbach, M. L. Psychedelia: The interplay of music and psychedelics. Ann. N. Y. Acad. Sci. 1531, 12–28 (2024).

37. Driscoll, L. N., Duncker, L. & Harvey, C. D. Representational drift: Emerging theories for continual learning and experimental future directions. Curr. Opin. Neurobiol. 76, 102609 (2022).

38. Gillet, S. N., Kato, H. K., Justen, M. A., Lai, M. & Isaacson, J. S. Fear learning regulates cortical sensory representations by suppressing habituation. Front. Neural Circuits 11, 1–9 (2018).

39. Chaloner, F. A. & Cooke, S. F. Multiple Mechanistically Distinct Timescales of Neocortical Plasticity Occur During Habituation. Front. Cell. Neurosci. 16, 1–15 (2022).

40. Brockett, A. T. & Francis, N. A. Psilocybin decreases neural responsiveness and increases functional connectivity while preserving pure-tone frequency selectivity in mouse auditory cortex. J. Neurophysiol. 45–53 (2024) doi:10.1152/jn.00124.2024.

41. Henschke, J. U., Price, A. T. & Pakan, J. M. P. Enhanced modulation of cell-type specific neuronal responses in mouse dorsal auditory field during locomotion. Cell Calcium 96, 102390 (2021).

42. Niell, C. M. & Stryker, M. P. Modulation of Visual Responses by Behavioral State in Mouse Visual Cortex. Neuron 65, 472–479 (2010).

43. Kelmendi, B., Kaye, A. P., Pittenger, C. & Kwan, A. C. Psychedelics. Curr. Biol. 32, R63–R67 (2022).

44. Nieto-Diego, J. & Malmierca, M. S. Topographic Distribution of Stimulus-Specific Adaptation across Auditory Cortical Fields in the Anesthetized Rat. PLoS Biol. 14, 1–30 (2016).

45. Duque, D., Pérez-González, D., Ayala, Y. A., Palmer, A. R. & Malmierca, M. S. Topographic distribution, frequency, and intensity dependence of stimulus-specific adaptation in the inferior colliculus of the rat. J. Neurosci. 32, 17762–17774 (2012).

46. Condon, C. D. & Weinberger, N. Habituation produces frequency-specific plasticity of receptive fields in the auditory cortex. Behav. Neurosci. 105, 416–430 (1991).

47. Priebe, N. J. & Ferster, D. Inhibition, Spike Threshold, and Stimulus Selectivity in Primary Visual Cortex. Neuron 57, 482–497 (2008).

48. Chambers, A. R., Aschauer, D. F., Eppler, J., Kaschube, M. & Rumpel, S. A stable sensory map emerges from a dynamic equilibrium of neurons with unstable tuning properties. Cereb. Cortex 33, 5597–5612 (2022).

49. Deitch, D., Rubin, A. & Ziv, Y. Representational drift in the mouse visual cortex. Curr. Biol. 31, 4327-4339.e6 (2021).

50. Adesnik, H., Bruns, W., Taniguchi, H., Huang, Z. J. & Scanziani, M. A Neural Circuit for Spatial Summation in Visual Cortex. Nature 490, 226–231 (2012).

51. Willins, D. L., Deutch, A. Y. & Roth, B. L. Serotonin 5-HT(2A) receptors are expressed on pyramidal cells and interneurons in the rat cortex. Synapse 27, 79–82 (1997).

52. Marek, G. J. & Aghajanian, G. K. LSD and the phenethylamine hallucinogen DOI are potent partial agonists at 5-HT2A receptors on interneurons in rat piriform cortex. J. Pharmacol. Exp. Ther. 278, 1373–1382 (1996).

53. Tang, Z. H. et al. The effects of serotonergic psychedelics in synaptic and intrinsic properties of neurons in layer II/III of the orbitofrontal cortex. Psychopharmacology (Berl). 240, 1275–1285 (2023).

54. de Filippo, R. et al. Somatostatin interneurons activated by 5-ht2a receptor suppress slow oscillations in medial entorhinal cortex. Elife 10, 1–21 (2021).

55. Cisneros-Franco, J. M., Ouellet, L., Kamal, B. & de Villers-Sidani, E. A Brain without Brakes: reduced Inhibition Is Associated with Enhanced but Dysregulated Plasticity in the Aged Rat Auditory Cortex. Eneuro 5, ENEURO.0051-18.2018 (2018).

56. Zheng, Q. Y., Johnson, K. R. & Erway, L. C. Assessment of hearing in 80 inbred strains of mice by ABR threshold analyses. Hear. Res. 130, 94–107 (1999).

57. Romero, S. et al. Cellular and Widefield Imaging of Sound Frequency Organization in Primary and Higher Order Fields of the Mouse Auditory Cortex. Cereb. Cortex 30, 1603–1622 (2020).

58. Pachitariu, M. et al. Suite2p: beyond 10,000 neurons with standard two-photon microscopy. bioRxiv. Bioarxiv 20, 2017 (2017).

59. Baden, T. et al. The functional diversity of retinal ganglion cells in the mouse. Nature 529, 345–350 (2016).

